# ***noisyR:*** Enhancing biological signal in sequencing datasets by characterising random technical noise

**DOI:** 10.1101/2021.01.17.427026

**Authors:** I. Moutsopoulos, L. Maischak, E. Lauzikaite, S. A. Vasquez Urbina, E. C. Williams, H. G. Drost, I. I. Mohorianu

## Abstract

High-throughput sequencing enables an unprecedented resolution in transcript quantification, at the cost of magnifying the impact of technical noise. The consistent reduction of random background noise to capture functionally meaningful biological signals is still challenging. Intrinsic sequencing variability introducing low-level expression variations can obscure patterns in downstream analyses.

We introduce ***noisyR***, a comprehensive noise filter to assess the variation in signal distribution and achieve an optimal information-consistency across replicates and samples; this selection also facilitates meaningful pattern recognition outside the background-noise range. ***noisyR*** is applicable to count matrices and sequencing data; it outputs sample-specific signal/noise thresholds and filtered expression matrices.

We exemplify the effects of minimising technical noise on several datasets, across various sequencing assays: coding, non-coding RNAs and interactions, at bulk and single-cell level. An immediate consequence of filtering out noise is the convergence of predictions (differential-expression calls, enrichment analyses and inference of gene regulatory networks) across different approaches.

**Teaser:** Noise removal from sequencing quantification improves the convergence of downstream tools and robustness of conclusions.

## Introduction

High-throughput sequencing (HTS) became a new standard in most life science studies yielding unprecedented insights into the complexity of biological processes. The increase in sequencing depth and number of samples, across both bulk and single cell experiments, facilitated a greater diversity in biological questions (*1*), at the same time allowing a higher sensitivity for the detection of perturbations in gene expression levels between samples (*2*). This increased accuracy greatly assists with the biological interpretation of results such as identification and characterisation of differential expression (DE) at tissue and cellular levels (*3*) or the inference and characterisation of gene regulatory networks (*4*). However, HTS may exhibit high background noise levels resulting from non-biological/technical variation, introduced at different stages of the RNA-seq library preparation, or from amplification/sequencing bias (*5*) to random hexamer priming during the sequencing reaction (*6*). These technical alterations of signal can affect the accuracy of the downstream DE results or create spurious patterns biasing downstream interpretations. Statistical methods developed to date (*7–9*), focused mainly on batch/background correction, normalisation, and evaluation of DE, have been designed to mitigate the impact of these biases on DE analyses (*10*). A noise filter for pre-processing of the data before these steps would ensure a reduction of further amplification of these biases. Here, we introduce a new high-throughput noise filter to remove random technical noise from sequencing data and illustrate the downstream information consistency that is achieved.

While technologies may exhibit different technical biases, the sequencing bias across an experiment was expected to be uniform. This expectation was based on the assumption that sequencing reads would uniformly cover the expressed transcripts, with the algebraic sum of reads from each gene being proportional to the expression of that gene (*11*). However, in practice we observe a reproducible, yet uneven distribution of signal across transcripts (*11*); moreover, highly abundant genes show a higher consistency of transcript-coverage than lower abundance genes. This coverage bias of lower abundance genes is one of the main origins of technical noise (*12*). The latter can be attributed to the stochasticity of the sequencing process, the limits of sequencing depth, and alignment inaccuracies during the mapping procedure. To further explore the coverage bias of lower abundance genes, we define genes whose quantification is characterised by such a lack of coverage-uniformity as “noisy”.

The presence of noise in HTS data has been widely acknowledged, and there have been several attempts to understand and quantify it. A recent study (*13*) presented a variety of common experimental errors that may increase sequencing noise and proposed ways to alleviate their effect such as using a mild acoustic shearing condition to minimise the occurrence of DNA damage. Fischer-Hwang and colleagues (*14*) presented a denoising tool that can be applied on aligned genomic data with high fold-coverage of the genome to improve variant calling performance. The recent prevalence of single-cell sequencing technologies has further highlighted the issue of noise, as the lower sequencing depth per cell leads to more uncertainty of the quantification of (low abundance) genes. Efforts have been made to reduce the noise levels experimentally, such as by utilising a different barcoding approach (*15*).

On the computational side, several imputation and denoising algorithms have been proposed, e.g. a machine learning (ML) based deep count autoencoder (*16*). Other tools focus on DE analysis, such as TASC (*17*), which uses a hierarchical mixture model of the biological variation. However, successful methods usually rely on assumptions specific to the biological experiment and are tailored to particular settings or model systems, thus leaving most large-scale sequencing efforts, lacking such specific experimental design, exposed to random technical noise. To our knowledge, there is little focus on bulk experiments, where technical noise still exists at low abundances, independent of biological assumptions; for these experiments the low number of replicates hinders imputation-based approaches.

Existing approaches for calling DE genes mitigate to various extents the presence of noise, however these are not designed to identify and assess the impact of genes showing random, low-level variation. As a result, some of these are detected by the DE analyses, biasing the biological interpretation of the results. In addition, the choice of tools used for preprocessing steps may influence the relative transcript expression estimation accuracy (*18*). These analytical biases mainly arise from differences in the detection and handling of transcript isoforms or processing of unmapped and multi-mapping reads (*3*). Such variation in abundance estimation can in turn strongly affect the downstream analyses (*19*).

We developed ***noisyR***, a denoising pipeline to quantify and exclude technical noise from downstream analyses, in a robust and data-driven way. The approach underlines consistency of signal over a user-defined threshold. ***noisyR*** is applicable on either the original, unnormalised count matrix, or alignment data (BAM format). Noise is quantified based on the correlation of expression across subsets of genes for the former, or distribution of signal across the transcripts for the latter, in different samples/replicates and across all gene abundances (Methods). We illustrate the approach on bulk and single cell RNA-seq datasets and highlight the impact of the noise removal on refining the biological interpretation of results.

## Results

### Noise quantification in bulk RNA-seq data

To exemplify the impact of denoising on the biological interpretations from bulk RNA-seq experiments, we applied ***noisyR*** on mRNA-seq and smallRNA-seq data. First, we illustrated the advantages of using the pipeline on a subset of mRNA-seq samples from a 2019 study by Yang and colleagues (*20*). To assess the distributions of signal we used density plots (Fig. 1A) and summaries of Jaccard similarity indices (Fig. 1B) across all samples. For the former, we observed a multi-modal distribution that suggests a signal to noise transition range between [3,7] on log2 scale; for the latter, the high similarity along the diagonal mirrors the temporal component of the time series. To reduce the number of low abundance, high fold change DE calls (Fig. 1C, fig.S1A for sample similarity and the secondary DE distribution visible in Fig. 1D and fig. S1C), we used first the ***noisyR*** count-based pipeline, on default parameters: window length = 10% x #genes and sliding step = 5% x window length (Fig. 1, E and H, fig. S1E). We used a correlation threshold of 0.25 and the boxplot median method, a combination of hyper-parameters producing the smallest coefficient of variation across abundance thresholds for the considered samples (Methods); the interquartile ranges (IQRs) of noise thresholds for the different samples ranged between 39 and 63, with an average of 58, for sequencing depths varying between 58M and 82M (Fig. 1E, fig. S1E). We detected an outlier with a low threshold of 18 (corresponding to a sequencing depth of ~77M) and three with values of over 100, corresponding to sequencing depths of 73M, 71M and 96M respectively. Next, we applied the transcript approach focusing on the correlation of the expression profiles across exons/transcripts (Methods); despite the higher runtime compared to the count-based approach, the transcript-approach was more robust, as illustrated by the lower variance in signal/noise thresholds across samples (Fig. 1I). The parameters that minimised the coefficient of variation were: correlation threshold = 0.26 and the boxplot median method; the resulting noise threshold IQRs ranged between 64 and 79, with an average of 75 and one outlier at 104. The signal/noise thresholds were similar for the two options, with an increased level of detail for the transcript-based approach.

**Fig. 1.**
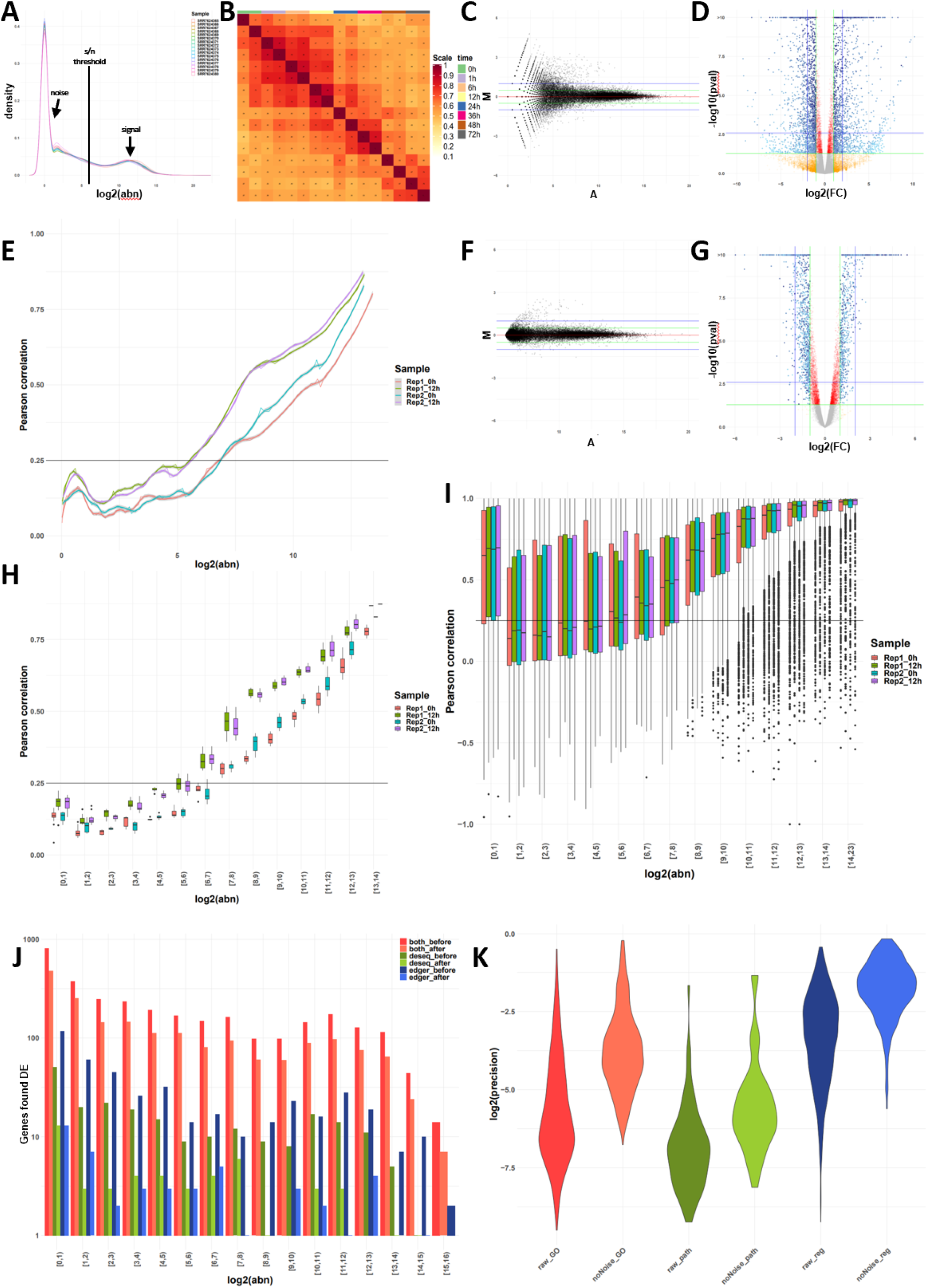
Overview of QC measures and original vs denoised outputs on standard components of an mRNA-seq pipeline. **(A)** Distributions of gene abundances by sample; the RHS distribution corresponds to the biological signal, the LHS distribution to the technical noise; the aim of ***noisyR*** is the identification of biologically meaningful values for the signal/noise threshold in between. **(B)** JSI on the 100 most abundant genes per sample; the replicates, and consecutive time points share a larger proportion of abundant genes. **(C)** MA plot of the raw abundances for the two 12h biological replicates; a larger proportion of low abundance genes exhibit high fold-changes, potentially biasing the DE calls. **(D)** Volcano plot of differentially expressed genes on the original, normalised count matrix; the colour gradient is proportional to the gene abundance. **(E)** Line plot of the PCC calculated on windows of increasing average abundance for the count matrix-based noise removal approach. **(F)** MA plot of the de-noised abundances for the two 12h biological replicates; the low-level variation is significantly reduced. **(G)** Volcano plot of differentially expressed genes on the denoised count matrix. **(H)** Box plot of the PCC binned by abundance for the count matrix-based noise removal approach. **(I)** Box plot of the PCC binned by abundance for the transcript-based noise removal approach. **(J)** Histogram of the differentially expressed genes found by applying DESeq and edgeR on the original and denoised count matrix respectively, binned by abundance; counts are on a logscale for visualization. **(K)** Violin plot of the precision (intersection size divided by the query size) for the results of the enrichment analysis performed on the differentially expressed genes found for the original (*raw*) and denoised (*noNoise*) matrices (log-scale). In the Gene Ontology set (*GO*) the terms from Biological Processed, Cellular Component and Molecular Function were grouped; in the Pathway set (*path*) the *Kegg* and *Reactome* terms were grouped; in the Regulatory terms (*reg*) the enriched Transcription Factors and microRNA entries were grouped.

These thresholds were used to exclude noisy genes from the count matrix (~44k genes were excluded out of ~56k genes expressed); the number of retained genes were 19.7k and 15.6k for the counts and transcript approaches, respectively. As a DE pre-processing step, the averaged noise threshold was added to all entries in the count matrix (Methods). The effect of the noise removal is illustrated by the narrower distribution in the MA plots (Fig. 1F, fig. S1B). Next, we performed a DE analysis between the 0h and 12h samples of the Yang dataset using the denoised matrix. Following the noise correction, we saw a 46% reduction in the number of DE genes – from 3,607 to 1,952. A large number of low abundance genes with spuriously high fold-changes were no longer called DE (*12*). Moreover, when comparing the outputs of two standard DE pipelines, edgeR (*8*) and DEseq2 (*7*), we noticed that the number of genes identified as DE by both methods only marginally decreased when the noise corrected input is used, whereas the number of DE genes called only with edgeR or only with DeSeq2 decreased significantly (Fig. 1J, fig. S1F); therefore we observed an increase in output consistency across methods when the noise filtered inputs were used. Moreover, the fold-changes and p-values of denoised genes correlated better and we no longer saw a large set of DE genes with (adjusted) p-values marginally below the DE threshold (Fig. 1D vs G. fig. S1C vs D). This step was followed by a functional enrichment analysis focusing on the DE genes, with the genes expressed (post filtering) as background set (*21*). The number of enriched terms was lower in the denoised data, 1,108 vs 4,671 in the original analysis; ~24% of the terms were retained and the terms found with the denoised dataset were approximately a subset of the ones found without the noise correction (~99.6% of terms found after denoising were also found prior to noise removal). In addition, the noise-correction terms corresponded to a higher percentage of genes assigned per pathway (Fig. 1K). Thus, applying ***noisyR*** focused the interpretation of results on the enrichment terms with highest confidence, ensuring biological relevance.

The ***noisyR*** transcript approach was also applied on two small RNA (sRNA) datasets, from plants (*A. thaliana*) and animals (*M. musculus*), respectively. In contrast to the mRNAseq data, sRNAs samples had different correlation vs abundance distributions. Overall low abundance sRNA transcripts/loci contained more noisy entries (*22*). Also, we observed a sharper increase to high correlation entries highlighting the transition from degraded transcripts to precisely excised sRNAs (*23, 24*). For both model organisms, miRNA hairpins and transposable elements (TEs) were analysed separately. For the former, we observed overall higher correlations than for mRNAs, likely because of the precise cleavage of the mature duplex and the lack of signal outside the duplex region (*25*); this characteristic is stronger for the animal case (fig. S2C). For both animals and plants, the increasing distribution was clearly detectable (fig. S2, A and C). The TE distributions also reflected the characteristics of the underlying sRNAs; for the animal example (fig. S2D) we saw a sharper increase along the abundance bins, specific for the piRNAs (*26*), whereas in plants (fig. S2B), the distribution of signal (expressed siRNAs) mirrored the biogenesis of heterochromatin siRNAs (*27*).

### Effect of noise on single cell (smartSeq) data

To illustrate the broad applicability of ***noisyR*** on different HTS data, we present its output on single cell (smartSeq2) sequencing output focusing on a subset of samples from the dataset presented by Cuomo and colleagues (*28*); we focused on 6 donors, and one timepoint, the number of cells per donor varied between 45 and 107. A common difficulty in single-cell experiments is that due to the higher number of samples/cells, the runtime is much higher if the pipeline is applied without modification, making the transcript approach intractable in practice, for higher number of cells; we also assessed whether the inferred signal/noise threshold was informative.

First, we applied ***noisyR*** using the count matrix approach on all cells with default parameters; we observed that correlation values rose to a weakly positive plateau (0.2-0.4) and remained stable over a wide range of abundances (Fig. 2A). Our interpretation of this result is that lower sequencing depths and higher resolution of smart-seq compared to bulk data induces more dissimilarity across medium abundance values. To alleviate this effect, we grouped cells into a small number of “pseudo-samples”, both randomly and according to the sample origin (i.e. donor). For each pseudo-sample, we applied the count-based approach on the averaged expression of genes In the resulting ***noisyR*** output, we observed a clearer step in the abundance-correlation plot (Fig. 2B, Fig. S3A), especially when the cells were grouped by donor. This indicates that an effect of the summarisation is a reduction in cell-to-cell variability which also focuses the noise identification procedure. The thresholds obtained via pseudo-sample summarisation and count-based noise identification varied between 2 and 4 with an average of 2.6 (corresponding to a sequencing depth per pseudo-sample between 590K and 689K, representative of the average sequencing depth per cell of 640K); these were used in a similar manner as for the bulk data, to produce a denoised count matrix.

**Fig. 2.**
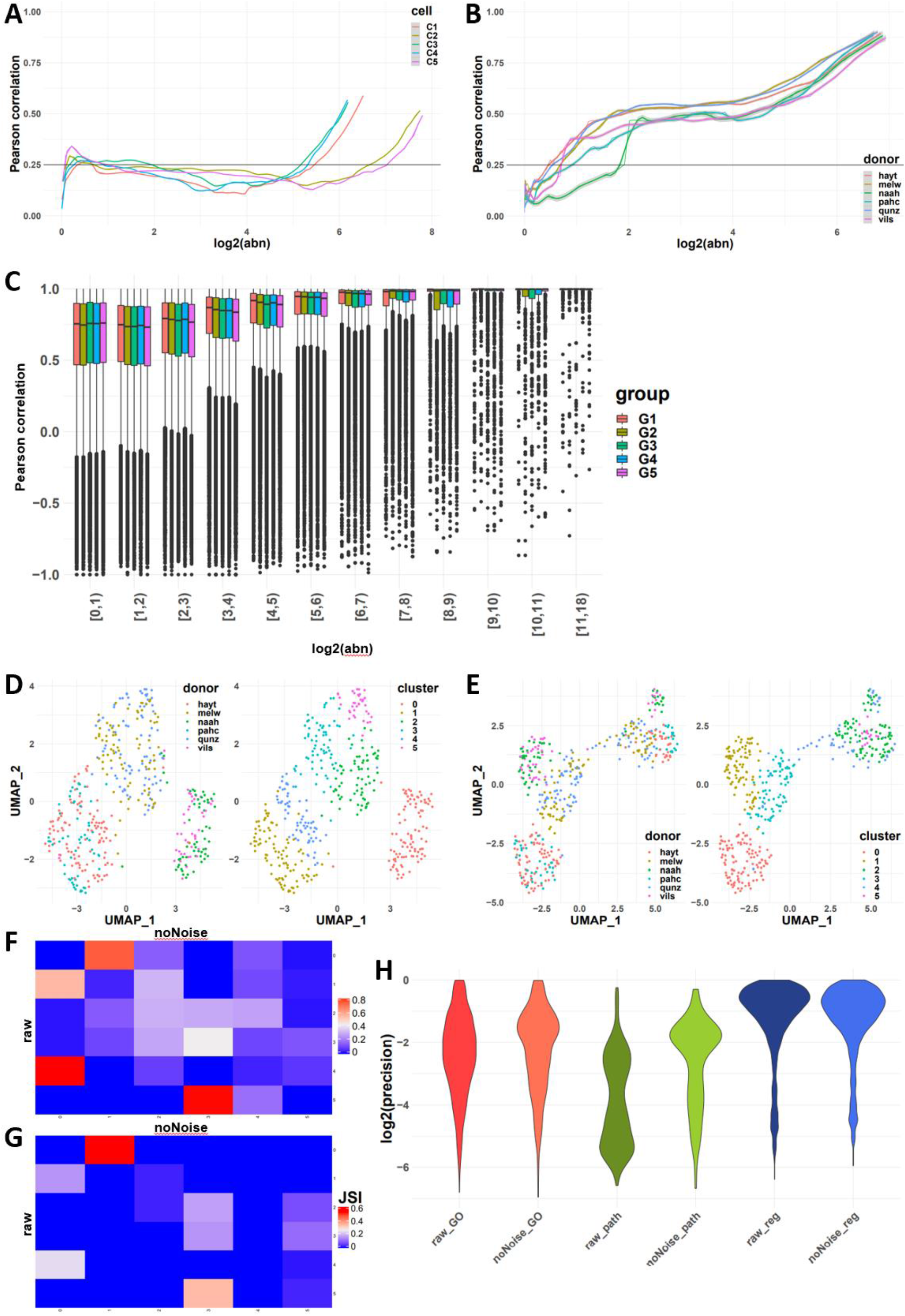
Overview of noise filtering on smartSeq data and impact on biological interpretation of results. **(A)** PCC calculated on windows of increasing average abundance for the count-matrix based noise removal approach applied to the full count matrix of all cells (four cells shown). **(B)** PCC calculated on windows of increasing average abundance for the count-matrix based noise removal approach applied to the “pseudo-samples” formed by grouping all cells from each donor. **(C)** Box plot of the PCC binned by abundance for the transcript-based noise removal approach applied to five groups of five cells each obtained by concatenating the corresponding BAM files. **(D)** UMAP representation of the cells using the raw count matrix grouped by donor (left) and by inferred cluster (right). **(E)** UMAP representation of the cells using the denoised count matrix grouped by donor (left) and by inferred cluster (right) **(F)** Contingency matrix of the clusters formed before and after the noise removal; the shade of each tile represents the proportion of the cluster from the raw matrix (row) that belongs to the corresponding cluster of the denoised matrix (column). **(G)** Heatmap of the Jaccard similarity index between the 50 most significant markers identified for each cluster on the raw matrix (rows) and denoised matrix (columns). **(H)** Violin plot of the precision (intersection size divided by the query size) for the results of the enrichment analysis performed on the marker genes found for each cluster of the raw and denoised matrix respectively (log-scale).

As the transcript approach is more computationally intensive, we applied it on a subsampled set of 25 cells. The subsamples were chosen randomly, and the process was reiterated five times, with the requirement that the summarised cells originate from the same donor. Formatting the data for ***noisyR*** was achieved by concatenating the BAM files for the selected cells and treating them as one sample. Whereas for the count approach, the results were highly variable between the cells, with several instances of low or negative correlations observed even at high abundances (Fig. 2A), the results obtained using the transcript approach with the concatenated BAM files were more consistent, with an expected increasing trend in the distribution of correlations (Fig. 2C). The correlation distributions were high, even at low abundances, which may be a consequence of the summarisation; a suitable threshold may be selected on the median, IQR, or 5-95% range to infer a signal to noise threshold, as the distributions are stable for low values and increase as the abundance increases above ~2 on a log2-scale.

To assess the impact of *noisyR* on the biological interpretation of results, we performed some downstream analyses before and after the noise removal and compared the results. In this study, we focus on the structure and mathematical characteristics of the outputs, rather than specific biological interpretations. The gene abundances were normalised and the cells were clustered using the Seurat R package (*29*) (see Methods). The different clusterings were visualised using the UMAP (non-linear) dimensionality reduction (*30*) (Fig. 2, D and E, fig. S3, B and C). We observed that cells clustered into three groups of two donors each when original data was used, suggesting a batch effect; However, cells corresponding to the four donors are mixed across clusters, when the denoised data was analysed, suggesting that some of the putative initial batch effect may had been alleviated with the noise correction. We also observed a better separation of clusters in the denoised data, especially on the first UMAP component, which may be an indication of robustness. We further assessed the similarity of the two clustering results using a cell-centred contingency table (Fig. 2F). We observed a good correspondence between the original and denoised matrices i.e. clusters 1 and 4 largely merged into cluster 0, and cluster 0 remains intact and turns into cluster 1. While the total number of clusters remained the same (under default parameters), the partitioning of cells was altered, which led us to believe that the results obtained with the original and denoised matrices may be qualitatively different, potentially affecting the downstream biological interpretations. To evaluate the changes in interpretation, we compared the clusters obtained prior to and post noise filtering by identifying the (positive) markers and computing the JSI between the top 50 markers of each cluster (Fig. 2G, fig. S3, D, E). Similarly as for the contingency table, the JSI heatmap shows an analogous correspondence between clusters, albeit weaker. Finally, we performed a functional enrichment analysis of the markers identified pre/post noise filtering. Similarly to the bulk results, there were fewer DE genes (markers per cluster) identified in the denoised dataset, with the precision being higher on average across the different GO terms, pathways, and regulatory terms (Fig. 2H). This strengthens our conclusion that the noise filtering process can add focus to the downstream biological analysis without significantly altering the overall composition of the data.

### Effects of noise filtering on the biological interpretation of regulatory interactions

One of the main aims of high-throughput sequencing projects, besides the identification of differentially expressed genes (the effect), is to infer the complex interactions of genes that lead to biological functions, the cause (e.g. disease, development or stress response). Understanding these interactions between genes and the corresponding regulatory elements (at transcriptional level, such as transcription factors (*31, 32*), or post-transcriptional, small RNAs (*33*)) allows us to unveil the molecular mechanisms encoding phenotypic outcomes, including causes of diseases.

#### Effect on PARE data on predicting regulatory miRNA/mRNA interactions

First, we sought to understand the effect of noise removal on the identification of miRNA/mRNA interactions. We applied the ***noisyR*** transcript approach to a Parallel Analysis of RNA Ends Sequencing (PAREseq) dataset (*34*). The distribution of degraded fragments across transcripts showed the same distribution of correlation vs abundance as we earlier observed for the bulk RNAseq data (Fig. 3A). Using a correlation threshold of 0.25, we determined a signal/noise threshold of 60 for this dataset. Next, we matched the highly abundant reads to known miRNAs (Methods, Fig. 3B) and illustrated that by removing the noisy reads, having abundance less than the noise threshold (Fig. 3, C-D), the prediction of interactions is simplified (*35*) i.e. for most genes only a few peaks were left. In some cases (e.g. Fig. 3C), only a very clear peak was retained after the noise removal, while for other transcripts some secondary interactions were kept. These results illustrate that noise-filtering is a crucial step for producing biologically meaningful mRNA/targeting predictions.

**Fig. 3.**
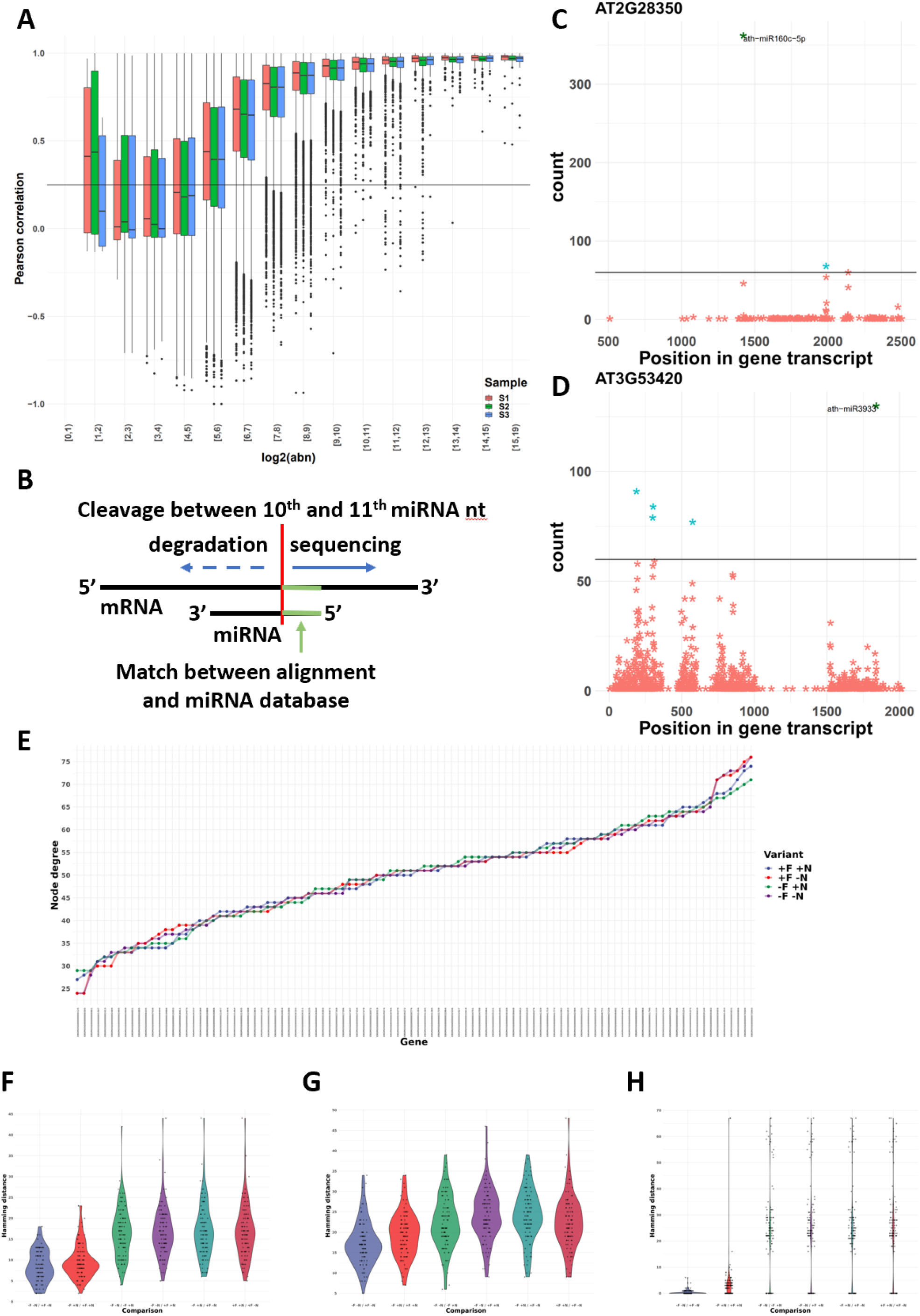
Effect of *noisyR* on PARE-Seq and GRN inference. **(A)** Box plot of the PCC binned by abundance for the transcript-based noise removal approach applied to PARE-Seq data. **(B)** Schematic overview of the microRNA/mRNA interaction; cleavage of the mRNA transcript occurs between the 10th and 11th nucleotide of the microRNA; **(C, D)** PARE t-plot illustrating the distribution of degradation products (each point) across the transcripts AT2G28350 and AT3G53420, respectively. All reads with summarised abundance less than the signal/noise thresholds are represented in red; degradation products corresponding to the signal, consistently identified across replicates, are represented in blue. The ones potentially generated by miRNAs are labelled. **(E)** node degree distributions (total number of edges connected to a node/gene) of 102 genes assigned to the neuron differentiation pathway from the Yang et al dataset. The four input data variants are shown: original (-F -N, purple); not noise-filtered but normalised (-F +N, green); noise-filtered but not normalised (+F -N, red); and noise-filtered and normalised (+F +N, blue) sorted by increasing values using -F -N as sorting key. (**F-H**) Pairwise hamming distance comparisons for each gene between all combinations of original (-F -N), noise-filtered (+F), and normalised (+N) input datasets using 102 Neuron differentiation genes from the bulk RNAseq (Yang et al.) dataset (Methods) show a comparable pattern across different gene regulatory network inference tools: **(F)** GENIE3; (**G**) GRNBoost2; (**H**) PIDC. The results consistently show that across network inference tools, noise-filtering has refining effects on the inferred network topologies in original or normalised data, further illustrating the advantages of noise-filtering to magnify biological signals by reducing technical noise.

#### Effect on the inference and interpretation of Gene Regulatory Networks

Characterising direct interactions between regulatory elements and their targets is only feasible for a limited set of interactions (such as the miRNA/mRNA interaction in plants, leading to mRNA degradation (*34*). To capture more of the vast complexity of gene interactions, for thousands of genes in tandem, Gene Regulatory Networks (GRNs) have been proposed as a systems biology tool to infer (direct and indirect) regulatory interactions from high-throughput sequencing data (expression data). In a gene regulatory network, nodes represent individual genes (e.g. transcription factors) and edges denote the regulatory interaction between connected genes. When edge-weights are considered, they encode the relative strength of the modeled interaction between two genes. After the network inference step, the resulting topology of GRNs can be used as a proxy for capturing the underlying biological and regulatory complexity of the studied process which in combination with enrichment analyses based on various Gene Ontologies generates a comprehensive model of the investigated process.

We evaluate the impact of noise-filtering on the inference of GRNs on particular network modules (subnetworks), associated with annotated pathways; we quantify the impact of random noise in altering network topologies and subsequent biological interpretations. To achieve this, we run our Network Inference Pipeline (NIP) and *edgynode* network analytics package (Methods) on bulk RNA-seq datasets using non-noise-filtered original, non-noise-filtered normalised, and noise-filtered normalised count matrices. Bulk RNAseq data has been widely used despite its well-known effect to dilute expression signals of individual cells or tissue types. However, in the context of technical noise, the averaging across cells and tissues may buffer the noise effect on general patterns while reducing the possibility to detect weak, but biologically meaningful, expression signals (e.g. transcription factor (*36*) or transposable element expression (*37*)).

Using the Yang dataset (*20*) in four different setups (original, -F(iltered) -N(normalised); noise-filtered but not normalised, +F -N; not filtered but normalised, -F +N; and noise-filtered and normalised, +F +N) and subsetted on five biological pathways (Placenta development, 46 genes; Neuron differentiation, 102 genes; Cell differentiation, 249 genes; Phosphorus metabolic process, 493 genes; and Multicellular organism development 996 genes), we ran NIP to infer GRNs using three inference approaches GENIE3, GRNBoost2, and PIDC, detailed in Methods. The inferred weighted correlation networks were imported into *edgynode* and rescaled to the range [0,100] to allow comparisons across inference tools.

Next, all rescaled weight matrices (fig. S4, E and F) were converted to binary format, using the median value over the entire weight matrix as threshold; a zero was assigned if the weight was below the median value, and a one, if the weight was above the median value. The resulting binary adjacency matrices were then used as input to compute the genespecific node degrees and to calculate the pairwise Hamming distances for each gene between combinations of original, noise-filtered, and normalised datasets (Fig. 3F, fig. S4, A-D) (Methods). This per-gene Hamming distance is a direct assessment of the number of edges that differ between inferences and captures both edge gain and loss. A low Hamming distance illustrates a robust network, whereas a high Hamming distance is proportional to large changes in the overall GRN topology. Panels Fig. 3, F-H illustrate pairwise comparisons between all combinations of input datasets: 1) original -F -N; 2) not noise-filtered but normalised -F +N; 3) noise-filtered but not normalised +F -N; and 4) noise-filtered and normalised +F +N exemplified for 102 genes corresponding to the neuron differentiation pathway and shown for all three network inference tools (GENIE3, Fig. 3F; GRNBoost2, Fig. 3G; and PIDC Fig. 3H). For all network inference tools, a common pattern is the refining effect of noise-filtering on the overall network topologies. Interestingly, the normalisation step has, in most cases, much greater impact on the network topology than noise-filtering. This result implies that the filtering procedure can detect and remove technical noise without disrupting the global network topology.

In addition, (fig. S4, E and F) shows a comparison between rescaled weight matrix distributions for an original and a noise-filtered and normalised network inferred with GENIE3. In this analysis, most genes had a large number of low-weight values within their edge-weight distributions that would result in thousands of biologically meaningless, weakly supported, connections with other genes. Noise-filtering in this bulk RNAseq dataset allows the exclusion of noisy genes as these fall below the median-threshold level which results in a more refined and biologically meaningful network topology after binarisation was applied (Methods).

Together, these results suggest that across network inference tools noise-filtering has refining effects on the inferred network topologies in original or normalised data, further illustrating the advantages of noise-filtering to magnify biological signals by reducing technical noise (*35*).

### *noisyR* package

The ***noisyR*** package is available on CRAN (https://CRAN.R-project.org/package=noisyr) and comprises an end-to-end pipeline for quantifying and removing technical noise from HTS datasets. The three main pipeline steps are [i] similarity calculation across samples, [ii] noise quantification, and [iii] noise removal; each step can be finely tuned using hyperparameters; optimal, data-driven values for these parameters are also determined. The package is written in the R (version 4.0.3) programming language and is actively maintained on https://github.com/Core-Bioinformatics/noisyR.

For the sample-similarity calculation, two approaches are available. The **count matrix approach** uses the original, un-normalised count matrix, as provided after alignment and feature quantification; each sample is processed individually, only the relative expressions across samples are compared. Relying on the hypothesis that the majority of genes are not DE, most of the evaluations are expected to point towards a high similarity across samples. Choosing from a collection of >45 similarity metrics (*38*), users can select a measure to assess the localised consistency in expression across samples (*12*). A sliding window approach is used to compare the similarity of ranks or abundances for the selected features between samples. The window length is a hyperparameter, which can be user-defined or inferred from the data (supplementary methods 1). The **transcript approach** uses as input the alignment files derived from read-mappers (in BAM format). For each sample and each exon, the point-to-point similarity of expression across the transcript is calculated across samples in a pairwise all-versus-all comparison. The output formats for the two approaches are the same; the number of entries varies, since the count approach focuses on windows, whereas for the transcript approach we calculate a similarity measure for each transcript.

The noise quantification step uses the abundance-correlation (or other similarity measure) relation calculated in **step i** to determine the noise threshold, representing the abundance level below which the gene expression is considered noisy e.g. if a correlation threshold is used as input then the corresponding abundance from a (smoothed) abundance-correlation line plot is selected as the noise threshold for each sample. The shape of the distribution can vary across experiments; we provide functionality for different thresholds and recommend the choice of the one that results in the lowest variance in the noise thresholds across samples. Options for smoothing, or summarising the observations in a box plot and selecting the minimum abundance for which the interquartile range (or median) is consistently above the correlation threshold are also available. Depending on the number of observations, we recommend using the smoothing with the count matrix approach, and the boxplot representation with the transcript option.

The third step uses the noise threshold calculated in **step ii** to remove noise from the count matrix (and/or BAM file). The count matrix can be calculated by exon or by gene; if the transcript approach is used, the exon approach is employed. Genes/exons whose expression is below the noise thresholds for every sample are removed from the count matrix. The average noise threshold is calculated and added to every entry in the count matrix. This ensures that the fold-changes observed by downstream analyses are not biased by low expression, while still preserving the structure and relative expression levels in the data. If downstream analysis does not involve the count matrix, the thresholds obtained in **step ii** can be used to inform further processing and potential exclusion of some genes/exons from the analysis.

## Discussion

### User-defined or data-driven options for the hyperparameters

***noisyR*** hyperparameters can be used to finely tune the identification of the signal/noise thresholds. To optimise the noise filtering procedure and dampen the stochastically induced differences between samples (e.g. derived from variation in sequencing depth or sample read-complexity) the noise removal step is performed by adding the average of the signal/noise thresholds across samples, on the raw count matrix. Nevertheless, comparable thresholds across the dataset are essential for a meaningful filtering; we recommend the use of consistency and robustness checks throughout the pipeline to ensure that the input samples are comparable, coupled with the data-driven selection of threshold values for setting hyper-parameters. The option of user-defined values is available, however the selected values should be based on observations from the input dataset, rather than exclusively following default recommendations. Next, we discuss in detail the options available for selecting the hyperparameters for a more adaptive noise-filtering based on the structure of the input data.

For the count matrix approach, the length of the sliding windows plays a significant role in assessing the similarity across samples. Smaller windows require more computational time; however the intended level of detail may not always be preferable, as small gene expression fluctuations, from sample to sample, would reduce the across-sample similarity if the abundance range is not wide enough (Fig. 5A). Even for medium-high abundances, expression or rank inconsistencies characterise smaller windows, indirectly leading to higher (and more variable across samples) signal/noise thresholds. If the window size is too large, less information is captured by the similarity measure and the accuracy of the noise threshold identification is also reduced (Fig. 5B). We recommend medium-sized windows that cover the abundance range in small incremental steps as larger overlaps between windows result in a more robust estimation of similarity-variation. An intuitive approach for determining an informative window size for a dataset relies on monotony changes of the similarity measure, quantified as the number of times the derivative of the correlation (as a function of abundance) changes sign. On several datasets, this resulted in a window length of 1/10th of the total number of expressed genes and a sliding window step size of 1/20th of the total gene number. A different tactic, also implemented in ***noisyR***, tackles this task from a different direction; it relies on optimising the window length using an entropy-based approach with the Jensen-Shannon divergence to assess the stability achieved as the window length is increased (supplementary methods 1). The shape of the distribution of correlations changes as the window length increases; however the change is less significant (evaluated using a t-test) for larger windows. The first point of stability is selected as the optimal window length, as it provides the largest possible granularity while maintaining robustness. The results from this approach are also consistent with earlier, empirical findings when applied to the Yang dataset (*20*).

**Fig. 4.**
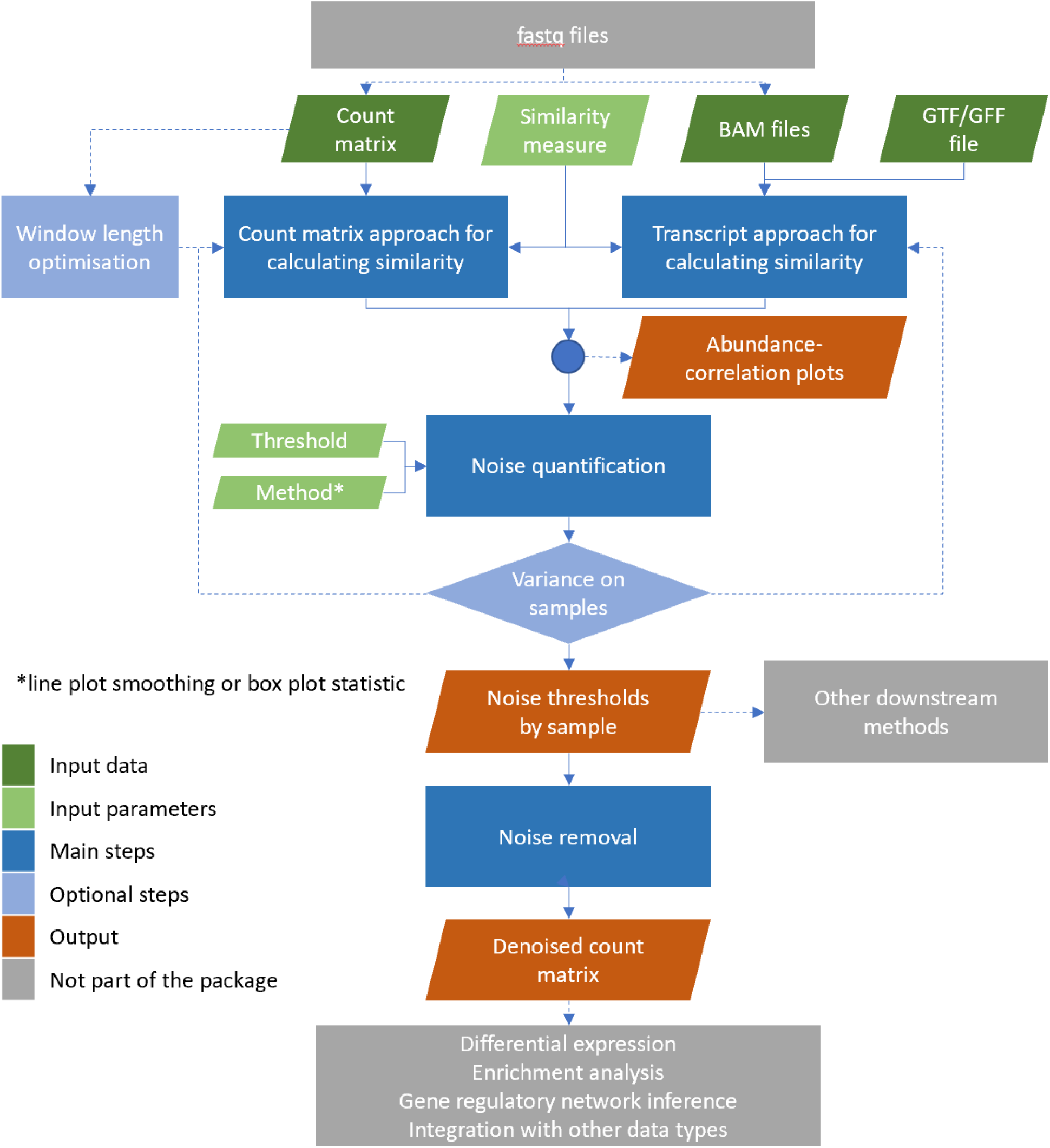
Workflow diagram of the noisyR pipeline. Workflow diagram describing the series of steps comprising the ***noisyR*** pipeline. Individual algorithms, finely tuned through hyper-parameters, are highlighted in blue. Optional steps are indicated through higher transparency. Common data pre- and post-processing steps not included in the package are indicated in grey.

**Fig. 5.**
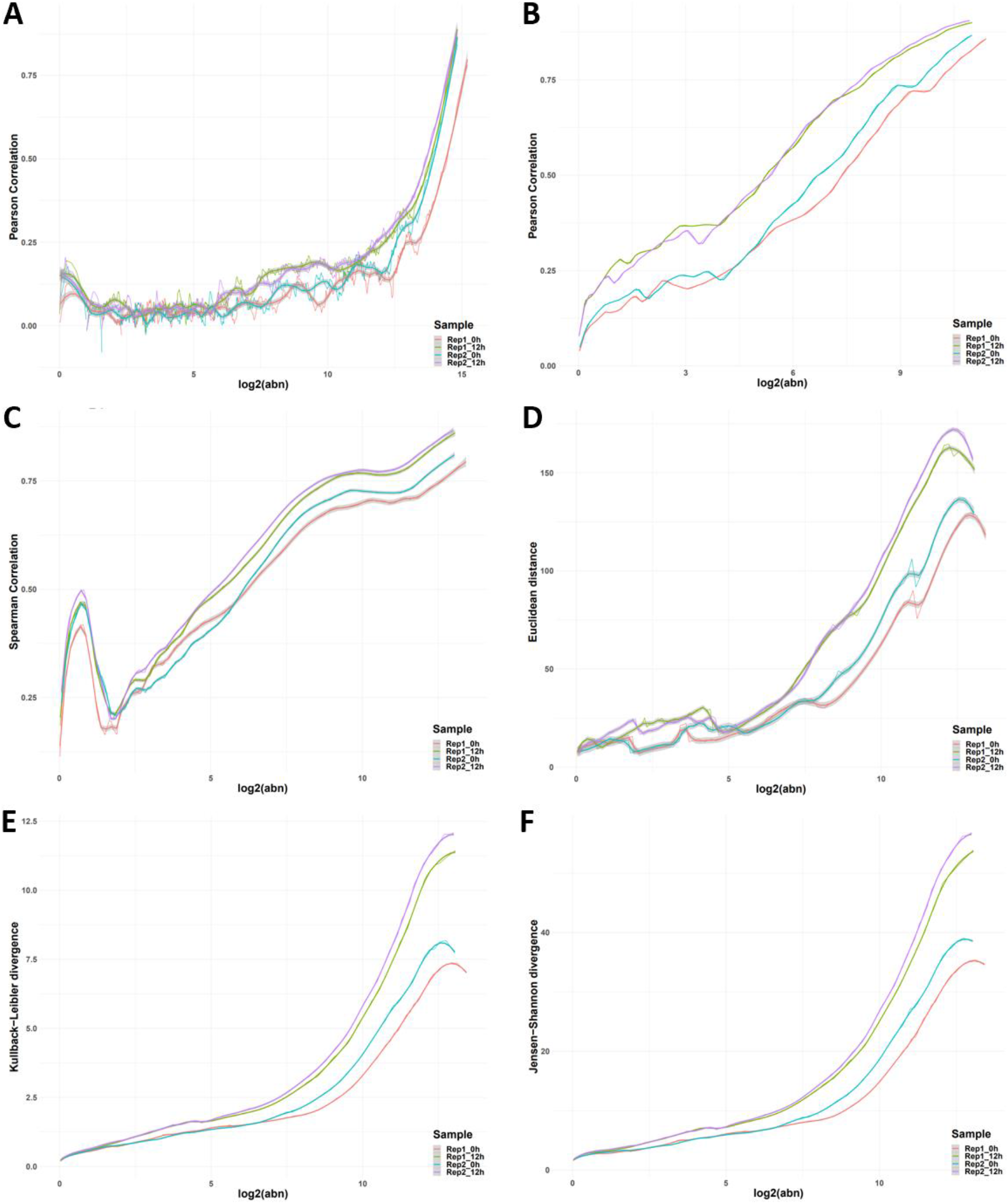
Effects of hyperparameter selection on noise quantification. **(A)** PCC-abundance plot for a window length of 1,000 genes, ~1/5th of the default. **(B)** PCC-abundance plot for a window length of 20,000 genes, ~4 times the default. **(C)** Spearman correlation plotted against abundance for the default window length of ~5,500. **(D)** Inverse of the Euclidean distance plotted against abundance for the default window length of ~5,500. **(E)** Inverse of the Kulbeck-Leibler divergence plotted against abundance for the default window length of ~5,500. **(F)** Inverse of the Jensen-Shannon divergence plotted against abundance for the default window length of ~5,500.

Yet another hyperparameter is the similarity measure; we compared the results for different correlation and distance metrics. We aim to achieve a high consistency in quantifying the signal/noise thresholds that is independent of the similarity measure. We tested the standard parametric and non-parametric correlation measures as well as the ones implemented in the *philentropy* package (*38*), which provides a variety of >45 distance measures. Dissimilarity measures are being inverted for comparison purposes (Fig. 5C-F illustrates the Spearman correlation, Euclidean distance, Kulbeck-Leibler divergence, and Jensen-Shannon divergence). Some measures have fixed ranges (e.g. the correlation coefficients), while others are semi- or unbounded. This raises the question of how to choose a similarity threshold when the range of values resulting from the similarity measure is unknown. Inspired by the correlation threshold, which provides a good separation at 0.25 for many datasets, we focus, as a starting point, on the naive assumption to use a quarter of the full range of the observed similarity values as a first cut-off approximation. Picking a threshold in a data-driven manner is, however, preferable and, in this case, achievable. Selecting from a variety of threshold values that minimise the coefficient of variation (standard deviation divided by the mean) of the corresponding noise thresholds in different samples is an empirical approach that works in practice. If the samples are semantically grouped e.g. replicates or time points, it may be better to minimise the variation in each individual group rather than across the full experimental design.

### Effect of aligner choice on noise quantification

The choice of the read-aligner was shown to influence the downstream DE analyses when the same quantification model was applied (*18*). To assess the effect of different alignment approaches on the quantification and observed levels of noise, mRNA quantification using featureCounts (*39*) was performed on reads aligned with STAR (*40*), HISAT2 (*41*) and Bowtie2 (*42*). The latter two were run both using their default parameters and with parameters set to match STAR functionality. For the count-based approach, the distribution of the Pearson Correlation Coefficients across abundance bins (Fig. 6A) shows that noise levels were relatively consistent regardless of the applied alignment algorithm. Similarly, for the transcript-based approach, the correlation distributions across abundance bins (Fig. 6B) illustrate little variation across aligners (additional examples in fig. S6, A and B). The estimated signal/noise thresholds were also comparable between the datasets generated by different aligners (Fig. 6C), with transcripts-based noise results being less variable. Once the noise correction was applied, the substantial peak in the abundance distributions around zero (Fig. 6D) was removed or significantly diminished and a second peak corresponding to the true signal was revealed around log2(abundance) of five using both counts and transcripts based approaches (Fig. 6,E and F respectively). The similarity of the abundance distributions across the datasets produced by the different aligners was observable both before and after the noise correction. This demonstrates that the proposed correction approaches are non-destructive and preserve the underlying biological signal. To further validate this point, the overlap between edgeR and DESeq2 analyses was investigated. The differentially expressed (DE) genes (adjusted p-value < 0.05 and |log2(FC)| > 1) detected by the two methods were compared for outputs produced using STAR (Fig. 1J), Bowtie2 (Fig. 6G) and HISAT2 (Fig. 6H). In all cases, there were fewer DE genes in total after noise correction was applied, and the specific differences for each DE method were reduced. The same conclusions were reached for the processing with Bowtie2 and HISAT2 applied with their default parameters (fig. S6C).

**Fig. 6.**
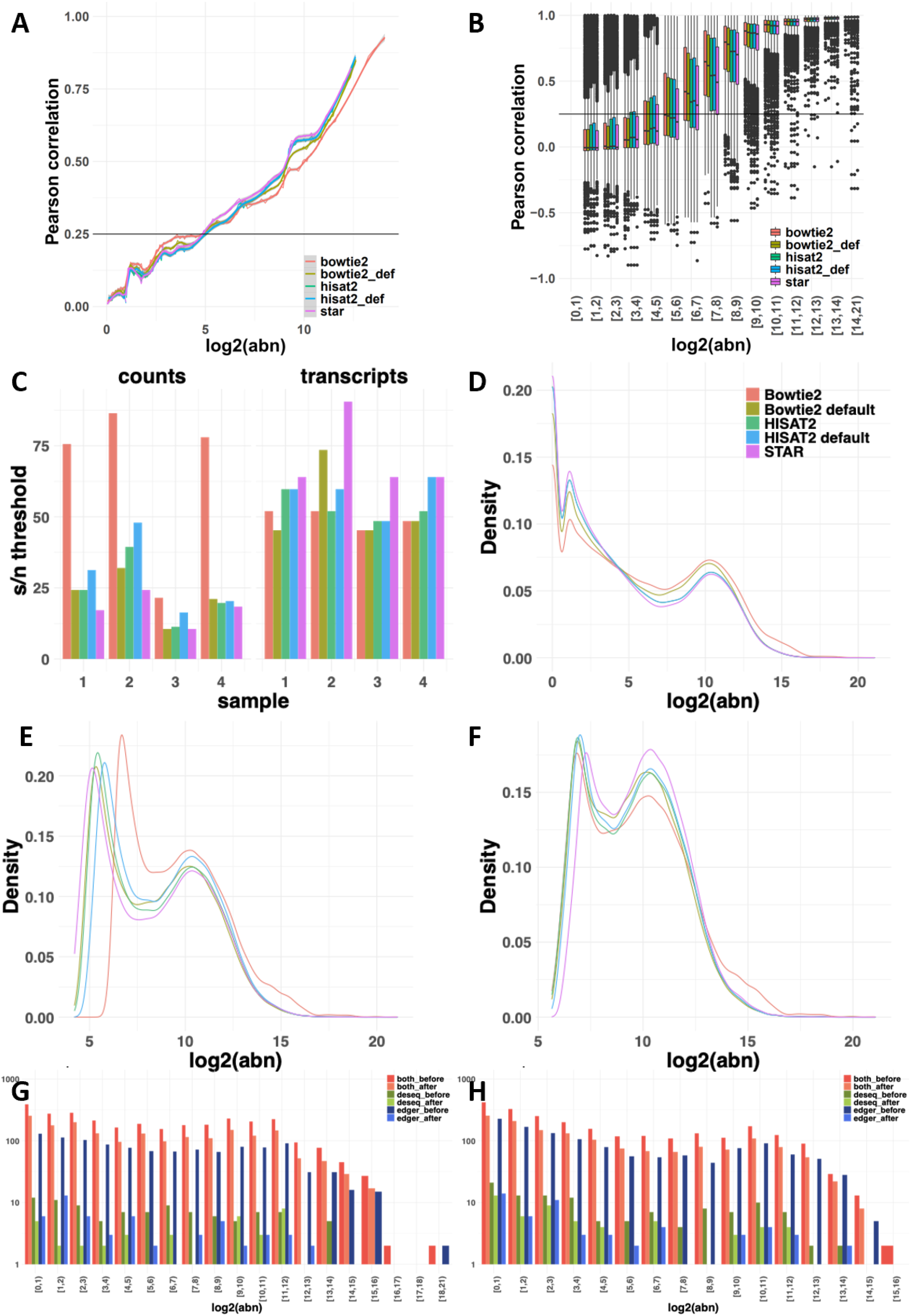
Assessment of aligner choice on noise quantification. **(A)** The distribution of PCC across abundance bins in datasets for a single mRNAseq sample obtained by STAR, Bowtie2 and HISAT2 alignment followed by featureCounts quantification using a counts-based noise removal approach. **(B)** The distribution of PCC across abundance bins in aligned read counts obtained by the five aligners for the same sample in the transcript-based noise correction approach. **(C)** The detected signal-to-noise thresholds in the four mRNAseq samples varied when the counts or transcripts-based noise correction methods were applied. **(D)** The distribution of abundance of reads aligned by the five algorithms and quantified by featureCounts. **(E)** The distribution of abundance of the quantified counts after counts-based noise correction **(F)** The distribution of abundance of the quantified counts after transcripts-based noise correction. **(G)** The number of the differentially expressed genes found by applying DESeq and edgeR on the original and denoised (using transcripts-based approach) count matrices obtained by Bowtie2 alignment. **(H)** The overlap between the DESeq and edgeR analyses performed on the original and denoised counts matrices obtained by HISAT2.

### The effects of noise-filtering on GRN inference for single cell RNAseq data

The recent emergence of single-cell sequencing technologies enabled the simultaneous assessment of expression variation between individual cells across thousands of celllineages. Although conceptually powerful, sequencing depths remain constrained by cost and in comparison to bulk RNAseq experiments the total number of reads is now shared among these (hundreds-) thousands of individual cells expressing thousands of genes each. This limit on the sequencing depth per cell underlines, yet again, the technical noise, whereby the quantification of low-abundance transcripts can be the result of either low biological expression or due to stochastic effects (likelihood) of read capturing. The requirement of an adaptive noise-filtering pipeline is fulfilled by ***noisyR***; the retained gene expression levels increases the robustness of quantification of single-cell data.

Analogous to the Yang et al. dataset, we used the Cuomo (*28*) dataset in four different setups (original, -F -N; noise-filtered but not normalised, +F -N; not filtered but normalised, -F +N; and noise-filtered and normalised, +F +N) and subsampled into three distinct biological pathways (Metabolism, 57 genes; Catalytic activity, 133 genes; Cellular metabolic process, 246 genes), we ran the Network Inference Pipeline to infer GRNs using the same three inference methods GENIE3, GRNBoost2, and PIDC (Methods) as used for bulk RNAseq data. The inferred weighted correlation networks were imported into *edgynode* and rescaled (fig. S5, C and D) analogous to the bulk RNAseq data shown in Results and Methods. The resulting pairwise Hamming distances for each gene between combinations of original, noise-filtered, and normalised datasets and for genes corresponding to various biological pathways (Fig. 7, A-C, fig. S5, A and B) show that total Hamming distances over all genes are larger in single-cell data. This implied that noise-filtering had a more significant/ refining impact on the inference and biological interpretations drawn from single-cell data when compared with analogous bulk RNA data.

**Fig. 7.**
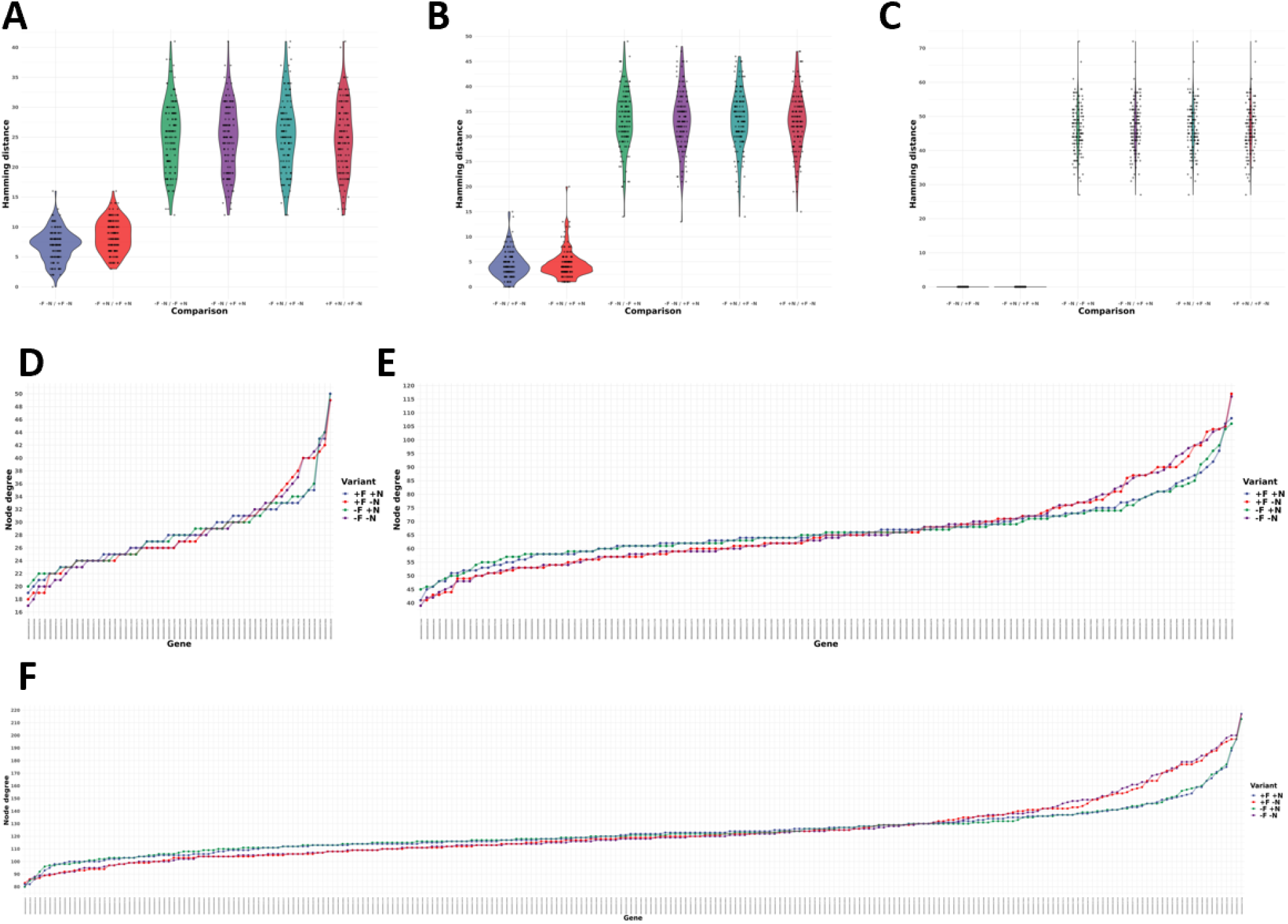
Node degree distributions and pairwise Hamming distance distributions between combinations of original, noise-filtered, and normalised input smartSeq datasets. **(A-C)** Pairwise Hamming distance comparisons for each gene between all combinations of original (-F -N), noise-filtered (+F), and normalised (+N) input datasets using 133 genes associated with catalytic activity pathways from the smartSeq (Cuomo et al.) dataset (Methods) show a comparable pattern across different gene regulatory network inference tools: **(A)** GENIE3; **(B)** GRNBoost2; **(C)** PIDC. The results consistently show that across network inference tools noise-filtering has refining effects on the inferred network topologies in original or normalised data, further illustrating the advantages of noise-filtering to magnify biological signals by reducing technical noise. **(D-F)**. For the smartSeq (Cuomoet al.) dataset the node degree distributions (total number of edges connected to a node/gene) of three sets of genes corresponding to different pathways are shown: **(D)** 57 genes associated with metabolism pathways; **(E)** 133 genes associated with catalytic activity pathways; **(F)** 246 genes associated with the cellular metabolic process. All four input data variants are shown: original (-F -N, purple); not noise-filtered but normalised (-F +N, green); noise-filtered but not normalised (+F -N, red); and noise-filtered and normalised (+F +N, blue) sorted by increasing values using -F -N as sorting key.

Fig. 7 D-F illustrates such analogous pairwise comparisons between all combinations of input datasets: 1) original -F -N; 2) not noise-filtered but normalised -F +N; 3) noise-filtered but not normalised +F -N; and 4) noise-filtered and normalised +F +N exemplified for 133 genes corresponding to catalytic activity pathways derived from single-cell RNAseq data (Cuomo et al.) and also shown for all three network inference tools (GENIE3, Fig. 7A; GRNBoost2, Fig. 7B; and PIDC Fig. 7C). Analogous to the bulk RNAseq results, noisefiltering has smaller effects on changes in network topologies than the normalisation step. Interestingly, it seems that the overall effect of noise-filtering in single-cell data has a stronger impact than in bulk RNAseq data (Fig. 3, F-H). Together, these conclusions hint toward a more useful effect of noise-filtering in single-cell data as is particularly expected for datasets with limited sequencing depth, but high individual cell numbers.

These highlight the positive effects of noise-filtering on magnifying meaningful biological signals in single-cell RNAseq data, with more significant effects in single-cell data due to the nature of technical noise induced by sequencing depth-constraints in combination with technical variation.

Using ***noisyR***, we demonstrate that unfiltered RNAseq quantifications can cause spurious false positive effects in various standard expression-based data analysis steps. To overcome these limitations, we introduce an R package to equip life scientists with a flexible solution, applicable across different bulk and single cell datasets, for excluding inconsistent transcript quantifications that would otherwise introduce stochastic variability in processed datasets. A comprehensive selection of automatic and semi-automatic threshold detection options provided by ***noisyR*** allows the robust inference of noise-thresholds to exclude low-confidence transcripts from processed RNAseq data. We illustrate the importance of such a noise-filtering procedure by assessing the convergence of DE identification and by inferring and comparing gene regulatory networks from various biological pathways, across gold-standard network inference tools. As a result, we find that noise-filtering is indeed able to significantly reduce stochastic effects magnifying underlying biological signals, thereby yielding more robust biological interpretations.

## Materials and Methods

### Materials

The bulk mRNA-seq used to illustrate ***noisyR*** was generated by Yang et al (*20*). The dataset comprises 16 samples across 8 time points [0-72 hours post stem cell induction. The raw data (fastq files and metadata) were downloaded from GEO (accession numbers GSE117896, GSM3314677 – GSM3314692).

Next, sRNA data was retrieved from Paicu et al (*43*) for the plant dataset (2 samples, a wildtype and DCL1 knockdown, with 3 biological replicates each, in *A. thaliana,* GSM2412286 – GSM2412291) and from Wallach et al (*44*) for the animal dataset, 6 samples generated for the identification of microRNAs as TLR-activating molecules in *M. musculus* (PMID: 31940779, GSE138532, GSM4110737 – GSM4110742). For both datasets, the reads were aligned to mature and hairpin miRNAs, downloaded from miRBase (*45*) and TEs, downloaded from TAIR and Ensembl, for *M. musculus.*

For assessing the impact of noise on direct biological interpretations and predictions, such as the interaction of miRNAs and mRNAs, we selected a PARE (parallel analysis of RNA ends, also known as degradome sequencing) dataset, consisting of 3 biological replicates (GSE113958) presented in Thody et al (*34*).

The single-cell mRNA-seq dataset used to illustrate ***noisyR*** was generated by Cuomo et al (study of stem cell differentiation) (*28*). The data is available on ENA, ERP016000 – PRJEB14362. The six donors with the highest number of cells (hayt, naah, vils, pahc, melw, qunz) were selected, cells in time point 3 were included.

The reference genomes used for alignment were: Homo_sapiens.GRCh38.98 (Ensembl version 98),: Mus_musculus.GRCm38.98 (Ensembl version 98) and *A. thaliana* (*46*).

### Methods, bulk mRNAseq data

#### Data pre-processing and quality checking

Initial quality checks were performed using fastQC (version 0.11.8), summarised with multiQC (version 1.9) (*47*). Alignments to reference genomes were performed using STAR (version 2.7.0a) with default parameters (*40*); the count matrices were generated using featureCounts (version 2.0.0) (*39*) against the *M. musculus* exon annotations obtained from the Ensembl database (genome assembly GRCm38.p6). Additional quality checks included density plots, (comparable distributions are a necessary but not sufficient condition for comparability), MA plots for the sufficiency check (expected to have a funnelling shape; observed outliers are candidates for differentially expressed transcripts), incremental dendrograms and PCA plots to evaluate the similarity of distributions (*12, 48*).

#### Data post-processing and biological interpretation of results

The differential expression analysis was performed after quantile normalisation of the count matrix using the standard functions from edgeR, version 3.28.0 (*8*) and DESeq2, version 1.26.0 (*7*). The thresholds for DE were |log2(FC)| > 1 and adjusted p-value < 0.05 (Benjamini-Hochberg multiple testing correction). The enrichment analysis was performed using g:profiler (R package gprofiler2, version 0.2.0) (*21*), against the standard GO terms, and the KEGG (*49*) and reactome (*50*) pathway databases. The observed set consisted of the DE genes, the background set comprised all expressed genes, using the full or de-noised count matrix respectively.

To assess the effect of noise correction across the multiple options of mRNA quantification, the sequencing reads were aligned to the reference genome using Bowtie2 (version 2.4.2) (*42*) and HISAT2 (version 2.1.0) (*41*). Aligners were run both with default parameters and with parameters set to match the STAR functionality of searching for up to 10 distinct, valid alignments for each read (“bowtie2 --end-to-end -k 10” and “hisat2 -q -k 10”). The transcript expression was quantified using featureCounts. The robustness of the quantification was assessed by investigating the overlap between edgeR and DESeq2 analyses. The genes with adjusted p-value < 0.05 (Benjamini-Hochberg multiple testing correction) and |log2(FC)| > 1 were considered before and after noise correction.

#### Gene regulatory network inference

To assess the implications of the noise filter on downstream biological interpretations, we used the bulk and single-cell datasets as inputs for various gene regulatory network (GRN) inference tools and compared the results for filtered and unfiltered inputs. For this purpose, we selected several gene subsets, ranging in size from 49 to 996 genes for the bulk dataset and from 57 to 246 genes for the single-cell dataset, based on enrichment analyses performed on the DE genes according to their inclusion in annotated pathways. (Supplementary table 1)

We chose a subset of the GRN inference tools benchmarked by BEELINE (*51*): GENIE3 (*52*), GRNBoost2 (*53*), and PIDC (*54*). We packaged the tools as Singularity containers (https://github.com/drostlab/network-inference-toolbox) and then assembled them into a custom pipeline (https://github.com/drostlab/network-inference-pipeline).

This pipeline extracts the subsets of genes corresponding to selected pathways and uses them as inputs for the GRN inference tools. The results are rescaled, binarised and compared using the *edgynode* package (v0.3.0, https://github.com/drostlab/edgynode). The edge weights and node degree distributions for all genes across the selected subsets are then visualised.

In detail, the similarity assessment of network topologies was performed using the *edgynode* function network_benchmark_noise_filtering() and was visualized using plot_network_benchmark_noise_filtering(). For this purpose, the inferred networks were converted to a binary format (presence/absence of an edge) using the overall median edge weight per network as a threshold. In network_benchmark_noise_filtering() four different types of matrices are used as input: a weighted adjacency matrix returned by a network inference tool where 1) no noise filter and no quantile normalisation (original) was performed (denoted in the figures as -F -N), 2) a noise filtering but no quantile normalisation was performed (+F -N), 3) no noise filtering but a quantile normalisation was performed (-F +N), and 4) both, noise-filtering and quantile normalization were performed (+F +N).

In a pairwise all versus all comparison, for each gene, the Hamming distance over the binary edge weight vectors was computed using the hamming.distance() function from the R package e1071 v1.7-4 (*55*), yielding a distribution of distances, which captures how many genes gained or lost their connection with other genes. A Kruskal-Wallis Rank Sum Test was performed using the stats::kruskal.test() function in R to assess whether comparisons of Hamming distance distributions between original, noise-filtered, and normalized combinations were statistically significantly different. Furthermore, visualising these distributions across comparisons and for all network inference tools facilitated an evaluation of the overall change of network topologies driven by the network inference tool or the normalisation/noise-filtering that was applied. These visualizations were then used to assess the impact and robustness of our noise-filter on the interpretation of biological network topologies. We applied the pipeline, including *edgynode*, with the same parameter configurations to both, bulk (Yang et al.) and single-cell (Cuomo et al.) data to retrieve comparable results for direct comparisons. Computationally reproducible analysis scripts to perform all inference steps, data transformations, and visualisations, including the ones used in this study can be found at https://github.com/drostlab/network-inference-pipeline.

### Methods, sRNAseq data

The 6 *A thaliana* sRNA samples were assessed using multiQC version 1.9 (*47*). Next, the sequencing adapters (both standard and HD) were trimmed using Cutadapt (version 3.2) (*56*) and the UEA sRNA Workbench (*57*). The larger 3 samples were subsampled without replacement to 8M reads (*12*); the smaller 3 samples were left unchanged. The read/sRNA-length distributions were bimodal with peaks at 21nt and 24nt, corresponding to miRNAs and TE-sRNAs, respectively. These sRNAs were aligned (using STAR (version 2.7.0a) (*40*)) to both microRNA hairpins (miRBase Release 22.1) (*45*) and TEs (obtained from TAIR10) (*46*).

The 6 *M. musculus* sRNA samples were processed in a similar way as the plant samples and subsampled without replacement to 3.5M sequences (*12*). The distribution of read lengths was bimodal with peaks at 22nt and 30nt corresponding to microRNAs and piRNAs respectively. The sRNAs were aligned to microRNA hairpins (miRBase Release 22.1) (*45*) and TEs (Ensembl release 101).

### Methods, PARE data

The 3 *A. thaliana* PARE samples (GSE113958) were QCed (multiQC version 1.9) (*47*) and the reads trimmed to 20nt; next, all samples were randomly subsampled without replacement to 25M (*12*). The subsampled reads were aligned to the reference genome (obtained from TAIR10 (*46*)) using STAR (using STAR (version 2.7.0a) (*40*)), with default parameters. The reads aligned to each position along a transcript were grouped on sequence and summarised by frequency. Each summarised fragment was matched (as reverse complement) to *A. thaliana* miRNAs. To visualise the distribution of signal across transcripts, t-plots were created, where each point corresponds to a summarised PARE fragment; the points for which a corresponding miRNA was identified were highlighted using the miRNA label (*34*).

### Methods, single cell data

For the single cell SmartSeq2 data, the cellranger software version 3.0 (*58*) was used for pre-processing, initial quality checks, and to generate the count matrix (it internally uses the STAR aligner). Further quality checks included distribution plots for the number of features, counts, mitochondrial and ribosomal reads per cell; significant outliers were removed during pre-processing. Dimensionality reduction and clustering were performed with the Seurat R package version 3.2 (*29*). The UMAP reduction method (*30*) was used for visualisation and assessment of results.

### Methods, noise quantification

Two approaches were implemented for the identification of noise. (1) The “count matrix approach” is a simple, fast way to obtain a threshold utilising solely the un-normalised count matrix (m genes x n samples). (2) The “transcript approach” is more refined, as it takes into account the distribution of signal across the transcript obtained by summarising the aligned reads from the BAM alignment files. For both approaches, a variety of correlation and distance measures are used to assess the stability of signal across samples (*38*). Most results were obtained using Pearson Correlation Coefficient (the default); similar results are obtained with other similarity or inverted dissimilarity measures such as Spearman Correlation, Euclidean distance, Kulbeck-Leibler divergence, and Jensen-Shannon divergence.

#### Count matrix approach

For each sample in the count matrix, the genes are sorted, in descending order, by abundance. A sliding window approach is used to scan the sorted genes (genes with similar abundances are grouped into “windows”). The window length is a hyper-parameter that can be user-defined or a single value inferred from the data using a Jensen-Shannon entropy based approach (supplementary methods 1). The sliding step can be varied to reduce computational time at the cost of reducing the number of data points and potentially losing accuracy. For each window, the correlation of the abundances of the genes from the sample of interest and all other samples is calculated and averaged using the arithmetic mean. Per sample, the variation in correlation coefficient (y-axis) is represented vs the average window abundance, x-axis. A correlation threshold (as a hyper-parameter) is used to determine a corresponding abundance threshold as a cut-off – the noise threshold. The correlation threshold is inferred from the data to minimise the variance of noise thresholds across the different samples. Several available approaches are based on the (smoothed) line plot or a binned boxplot of abundance against correlation (supplementary methods 2). Genes with abundances below the sample specific noise thresholds across samples were excluded from downstream analyses; the average of the thresholds were added to the count matrix, to avoid further biases. By increasing the minimum values in the count matrix from zero to the noise threshold, methods that are based on fold-changes will not emphasise small differences in abundance at very low values, which becomes especially problematic for genes that are seemingly absent in some samples but present and lowly expressed in others. This effect is particularly striking in single-cell data.

#### Transcript approach

Using the transcript coordinates of the aligned reads as input, the expression profile for each individual transcript was built as an algebraic point sum of the abundances of reads incident to any given position (*59*); if the alignment was performed per read, the corresponding abundance for every entry was set to +1. For each sample j, and for each transcript T, the point-to-point Pearson Correlation between the expression profile in j and the one in all other samples is calculated. The noise detection is based on the relative location of the distribution of the point-to-point Pearson Correlation Coefficient (p2pPCC) versus the abundances of genes and is specific for each individual sample. For low abundance transcripts the stochastic distribution of reads across the transcript leads to a low p2pPCC; the aim of the approach is to determine the range where the distribution of correlation coefficients (used as proxy for the distribution of reads across a transcript) are above a user-defined threshold; to approximate the signal-to-noise threshold a binning on the abundances was performed. For all examples presented in this study, the binning was done on log2 ranges; the signal-to-noise thresholds were defined as the abundance above which the first quartile of the p2pPCC distribution consistently remains above 0.25 (IQR method – see supplementary methods 2). Once a noise threshold was determined for each sample, the original count matrix was then filtered analogous to the count matrix approach. The BAM files can also be filtered directly by removing all genes which fall below the noise threshold in every sample. Downstream analysis that is not based on the count matrix, such as alternative splicing analysis can also be informed by the noise threshold by setting a lower bound of expression acceptance.

## Acknowledgments

IM, EL, EW and IIM acknowledge the constructive feedback from the Core Bioinformatics group and the support by the Core grant awarded to the Wellcome-MRC Cambridge Stem Cells Institute. SAVU, LM and HGD acknowledge that their work was supported by the Max Planck Society. IM and IIM designed the study and implemented the R package ***noisyR***; SAVU, LM and HGD implemented the R package *edgynode,* the analyses were performed by IM, IIM (the mRNA bulk and single cell analyses), EW, IIM (sRNA analysis), LM and HGD (GRN inference), EL, IM, IIM (comparison across tools). IM, HGD and IIM wrote the manuscript; all authors read and approved the submitted manuscript.

